# Simulating Short-Term Synaptic Plasticity on SpiNNaker Neuromorphic Hardware

**DOI:** 10.1101/2022.09.13.507796

**Authors:** Loïc J. Azzalini, Milad Lankarany

## Abstract

Neuromorphic chips are well-suited for the exploration of neuronal dynamics in (near) real-time. In order to port existing research onto these chips, relevant models of neuronal and synaptic dynamics first need to be supported by their respective development environments and validated against existing simulator backends. At the time of writing, support for short-term synaptic plasticity on neuromorphic hardware is scarce. This technical paper proposes an implementation of dynamic synapses for the SpiNNaker development environment based on the popular synaptic plasticity model by Tsodyks and Markram (TM). This extension is undertaken in the context of existing research on neuromodulation and the study of deep brain stimulation (DBS) effects on singular-neuron responses. The implementation of the TM synapse is first detailed and then, simulated for various response types. Its role in studies of DBS effect on postsynaptic responses is also reviewed. Finally, given the real-time capabilities offered by the hardware, we provide some insight to lay the groundwork for future explorations of closed-loop DBS on neuromorphic chips.

## 1 Introduction

By drawing inspiration from the organization of neurons in the brain, neuromorphic chips offer significant computational advantages, both in terms of speed and efficiency. These chips can be used to solve various real-world problems, which we regroup in the following categories: (i) AI and deep learning,

(ii) neuroscience discovery, and (iii) biomimetic sensors. Since these categories are not necessarily mutually exclusive, advances in one can be used for the advancement of another. For example, advances in the use of neuromorphic chips for neuroscience discovery can lead to the development of biologically-inspired algorithms which, in turn, can be utilized in AI and deep learning applications. Although the above categories might not cover all the benefits of neuromorphic technology, they can help us narrow down important features of different neuromorphic chips and how these may be capitalized for specific applications (Davies et al., 2021).

A key feature of neuromorphic chips utilized for neuroscience discovery is the ability to implement biophysical constraints and details towards the simulation of large neuronal networks. Initiatives such as the Human Brain Project (Markram et al., 2011), which include the SpiNNaker(S.B. Furber, Galluppi, Temple, & Plana, 2014) and BrainScaleS (Meier, 2015) neuromorphic systems, have accelerated computational neuroscience by allowing large-scale neuronal networks to be simulated (Sen-Bhattacharya et al., 2018). These systems are designed with scalability in mind, such that biophysical constraints introduced at the singular level may easily extend to larger ensembles.

By starting at the singular-neuron-level in this paper, we lay the groundwork for future explorations of real-time control systems with neuromorphic hardware in the context of neuroscience discovery. Since neuromorphic chips can run biological models in real-time (or near real-time), these chips can be used in model-based control applications, a control design that has garnered particular interest among up-and-coming closed-loop neuromodulation approaches (Lozano et al., 2019). Dedicated hardware allows models of brain function and dysfunction to be placed in the physical world where environmental effects and interactions may be investigated in the framework of closed-loop neuroscience. The exploration of deep brain stimulation (DBS) effects lends itself well to such brain-in-the-loop simulation.

A recent study (Milosevic et al., 2021) on the use of DBS for patient with movement disorders focused on the implication of short-term synaptic plasticity (STP) in understanding DBS effectiveness. Unlike long-term potentiation (respectively depression) which relates to the strengthening (weakening) of synapses on the order of seconds to minutes, STP describes transient dynamic effects that occur shortly after incoming spikes are received at the synapse of a neuron (on the order of milliseconds). There exist many models that describe the mechanics of neurotransmitter release and calcium influx responsible for these local effects (Barroso-Flores, Herrera-Valdez, Galarraga, & Bargas, 2017; Hennig, 2013), from the biophysical to the phenomenological (Markram, Wang, & Tsodyks, 1998). Given that DBS impact varies across brain regions and frequencies of stimulation, the authors suggest STP may account for the frequency dependence. While the study was carried out using site-specific responses for single neurons, extensions of the work, both in the scale of the simulation and the variety of the responses considered, would provide valuable insight to support the hypothesized role of STP.

Neuromorphic platforms such as SpiNNaker offer means to scale simulations from single neurons up to microcircuits effortlessly and as such, they are well suited to support this type of exploratory work. However, the simulation of DBS effects can only be carried out on neuromorphic hardware insofar as the models under study are supported by the platforms’s computational environment. Platform-agnostic programming interfaces such as pyNN (Davison et al., 2009) promote code reusability and validation, allowing neuron models to run on a workstation (e.g., through NEST (Gewaltig & Diesmann, 2007)) just as well as on neuromorphic hardware (e.g., SpiNNaker through its dedicated front-end sPyNNaker Rhodes et al. (2018)). Yet, varying levels of support across simulator backends limit portability. This work details steps taken to address this issue for the Tsodyks-Markram model of short-term synaptic plasticity. In doing so, we hope to contribute to the growing effort of cross-platform code sharing in computational neuroscience.

At the time of writing, the selection of synapse models on SpiNNaker is limited to static synapses (i.e., a constant weight) and various rules that fall under spike-timing dependent plasticity (STDP) dynamics. Furthermore, synapses projecting to the same postsynaptic target are required, by design, to be of the same type. In order to port our investigation of DBS effects (Milosevic et al., 2021) onto SpiNNaker, we need to extend existing functionality such that multiple input sources may be connected to the same postsynaptic neurons (or population thereof) via STP synapses. To this end, we propose a modification of the Python and C source code (accessible via the open-source sPyNNaker project (Rhodes et al., 2018)) that serves as the intermediary between PyNN (high-level) and the SpiNNaker hardware (low-level). Figure 1 provides an overview of the extended software functionality (in red) and it will be referenced in the following sections.

**Figure 1.**
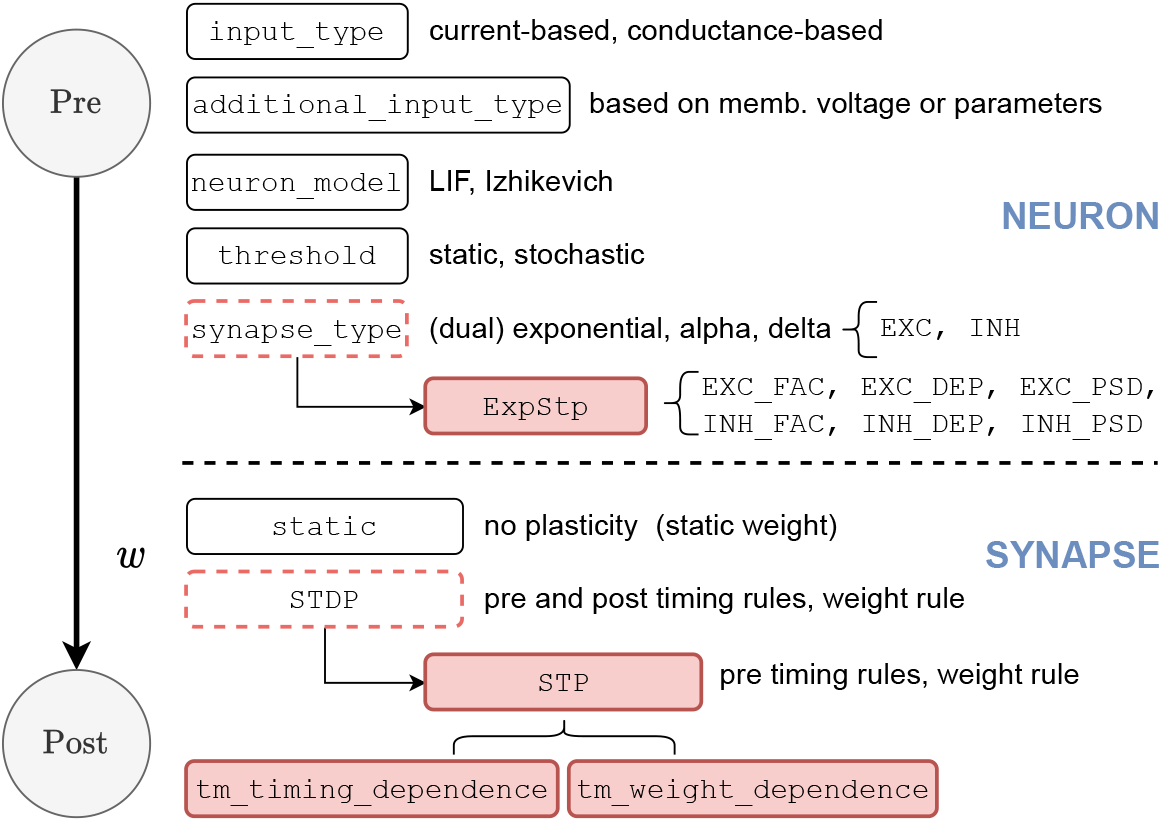
sPyNNaker neuron and synapse anatomy: the proposed STP implementation (shown in red) is derived from existing functionality

Section 2 introduces the Tsodyks-Markram model of short-term synaptic plasticity, alongside a roadmap detailing its implementation on SpiNNaker. The selection of a suitable TM model is achieved by considering existing implementations of STP on other platforms available through pyNN (Davison et al., 2009), the like of which may also be used as a means to validate our results. To this end, we also revisit the steady-state synaptic resources model proposed in (Markram et al., 1998). The second contribution of this work, which concerns the simulation of DBS effects on SpiNNaker, is introduced next. Finally, a generic leaky integrate-and-fire (LIF) neuron model is presented given its use in later simulations.

Simulation results for single and multiple STP synapses are presented in Section 3. The demonstration of facilitation and depression effects for a single TM synapse is followed by a comparison of postsynaptic effects on the LIF membrane voltage between the SpiNNaker and NEST implementations of STP. As an application of the new STP functionality on SpiNNaker, we present results depicting the simulation of DBS effects at different frequencies of stimulation.

The limitations of our implementation of STP on SpiNNaker are discussed in Section 4. In particular, concerns related to the scalability and the variability of the current TM synapse implementation are brought up; potential solutions in future extensions of this work are provided. Finally, additional applications motivating the use of STP on SpiNNaker within and without computational neuroscience are discussed.

## 2 Method

The models considered in this work are introduced below in order of perceived contribution. Each model presentation is followed by a walkthrough of their SpiNNaker implementation. Low-level implementations are written in C, while their corresponding PyNN interfaces are written in Python. To ensure correct compilation and integration of the resulting implementations with the sPyNNaker framework (to ultimately test these models on SpiNNaker), we use a template repository provided by the development team and made publicly available on GitHub (https://github.com/SpiNNakerManchester/sPyNNaker8NewModelTemplate).

### 2.1 Tsodyks-Markram Synapses

Following Milosevic et al. (2021), the implementation of short-term synaptic plasticity (STP) in this work is based on the popular phenomenological Tsodyks-Markram (TM) model of synaptic efficacy (Markram et al., 1998). By assuming limited amounts of resources at each synapse, the model captures dynamic effects related to neurotransmitter depletion (short-term depression) and those related to the influx of calcium at the presynaptic terminal (shortterm facilitation), in a manner that reflects the history of presynaptic activity (M. Tsodyks & Wu, 2013). Depression is modelled by the normalized variable *R* ∈ [0, 1], which denotes the amount of neurotransmitter resources available after synaptic transmission. On the other hand, the normalized utilization parameter *u* ∈ [0, 1] is used to denote the fraction of neurotransmitter resources ready for use, thus describing the competing facilitation effect (Farokhniaee & McIntyre, 2019). This description of synaptic efficacy has been used in many variants of the TM model over the years (Maass & Markram, 2002; Markram et al., 1998; M.V. Tsodyks & Markram, 1997). For reasons that will be made clear in the implementation walkthrough, the recursive TM equations proposed by Maass and Markram (2002) are used:

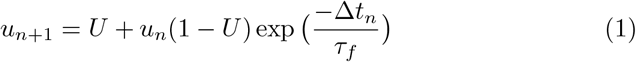

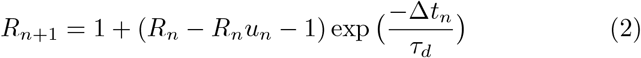

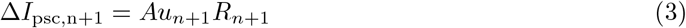

where *u*_*n*_ and *R*_*n*_ represent the state of the synapse immediately before the arrival of the *n*th spike; in turn, Δ*t*_*n*_ denotes the time elapsed between the (*n*+ 1)th and *n*th incoming spike. The increment in utilization parameter produced by an incoming spike is denoted by *U*, whereas *τ*_*f*_ is the time constant with which *u* decays to zero between spikes. On the other hand, *R* exponentially recovers with time constant *τ*_*d*_. The postsynaptic current (psc) input generated at the synapse in response to incoming spike is Δ*I*_psc,n_, where *A* denotes the absolute response amplitude corresponding to *u*_*n*_ = *R*_*n*_ = 1.0; Δ*I*_psc_ is referenced later in the text in relation to the synapse efficacy update.

We distinguish between three types of STP dynamic based on the selection of STP parameters *τ*_*f*_, *τ*_*d*_ and *U*. Based on a high-level interpretation of (1) and (2), short-term depression (D) is characterized by *τ*_*d*_ *>> τ*_*f*_, with large *U* (e.g., *U* = 0.5); conversely, *τ*_*f*_ *>> τ*_*d*_, with small *U* (e.g., *U* = 0.1), characterizes short-term facilitation (F). Additionally, synapses that exhibit a combination of depression and facilitation effects are described as being pseudo-linear (P) Farokhniaee and McIntyre (2019); Milosevic et al. (2021). These parameters may be selected based on observations of neuronal responses *in vivo*.

Given our focus on DBS effects later in the text, we are particularly interested in the steady-state behaviour of synaptic resource variables *u*_*n*_ and *R*_*n*_ when a TM synapse is subject to input spike trains of constant frequency *f*_DBS_. Theoretical knowledge of the long-term behaviour of a particular TM synapse also serves as a useful means of validating our implementation on SpiNNaker. We consider the steady-state resource variables *u*_*∞*_ and *R*_*∞*_ proposed by Markram et al. (1998) and recently revisited in (Ghadimi, Steiner, Popovic, Milosevic, & Lankarany, 2021):

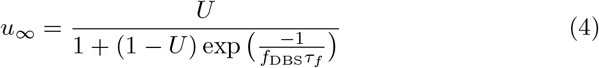

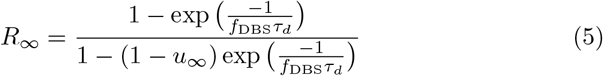

from which we derive the peak of the postsynaptic current at steady-state

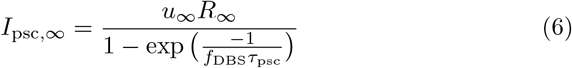

where *τ*_psc_ denotes the time constant of the postsynaptic exponential kernel which generates *I*_psc,n_.

#### Implementation

While STP is not supported by sPyNNaker *per se*, the existing STDP functionality (which is based on an “STDP core implementation”) may be reused to frame its implementation. Specifically, any feature relating to “post-before-pre” spike events may be disabled as STP dynamics only depend on the history of presynaptic spikes, that is the causal “pre-before-post” relationship between incoming spikes. In this framework, the synaptic rules are broken down into a timing-dependence and a weight-dependence (see “SYNAPSE” section in Figure 1). In order to be cross-compiled to run on SpiNNaker, the rules are implemented in C, while mirroring PyNN interfaces are written in Python.

The timing-dependence code is executed in response to presynaptic events (i.e., incoming spikes), allowing synaptic resource variables *u*_*n*_ and *R*_*n*_ to be updated according to (1) and (2) respectively. Given the low-level C environment, fixed-point arithmetic is used to implement the discrete-time equations, according to the time elapsed between the two most-recent spikes. More importantly, because divide and exponential operations are expensive on the SpiNNaker hardware (S. Furber & Bogdan, 2020), the facilitation and depression exponential functions are to be passed as look-up tables (LUT) after being precomputed in the mirroring, user-facing Python interface.

Given that the TM model phenomenologically describes a dynamic synapse, its implementation in simulation relies on the concept of synaptic weight *w*^*ij*^ (that which is kept constant in a static synapse), between a presynaptic neuron *j* and a postsynaptic neuron *i*. The weight-dependence code is responsible for updating the synapse weight *w*^*ij*^ after each presynaptic event to reflect strengthening (facilitation) or weakening (depression) of the synapse. As a result, the weight-dependence rule needs to be referenced in the timing-dependence code. Starting from a static weight 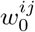 which, given multiple STP synapses to the same target, statically weighs their contribution, the TM synapse weight evolves according to

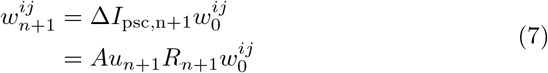

where typically *A* = 1.0, *u*_*n*+1_ and *R*_*n*+1_ are computed according to (1) and (2) respectively, and 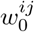 is set by the user during initialization.

In order to capture the STP dynamics of the TM model, it is important to observe the correct order of operations between the timingand weightdependence rules. For further details, we refer the reader to our implementation of the TM model for SpiNNaker (see Section 4) as well as the corresponding NEST implementation Gewaltig and Diesmann (2007). Given that the latter may also be accessed via PyNN, it is considered in Section 3 to assess the accuracy of our implementation.

### 2.2 DBS Effect

The addition of STP functionality to the SpiNNaker environment directly supports existing computational work on DBS effects Farokhniaee and McIntyre (2019); Milosevic et al. (2021), offering new directions in which to extend them. To simulate the effect of a single pulse of stimulation on a target neuron’s response, it is assumed that all presynaptic inputs are simultaneously activated (Milosevic et al., 2021). We consider a generic scenario illustrating this hypothesized DBS effect in Figure 2, where both excitatory and inhbitory presynaptic inputs project to the target via TM synapses (all three aforementioned types). The target is labelled as STN, for the subthalamic nucleus, which is a nucleus in the basal ganglia often selected as the stimulation site in the DBS treatment of Parkinson’s disease (Lozano & Lipsman, 2013). While only the response of a single STN neuron is considered in this work, it may represent a population of similar neurons in future extensions.

**Figure 2.**
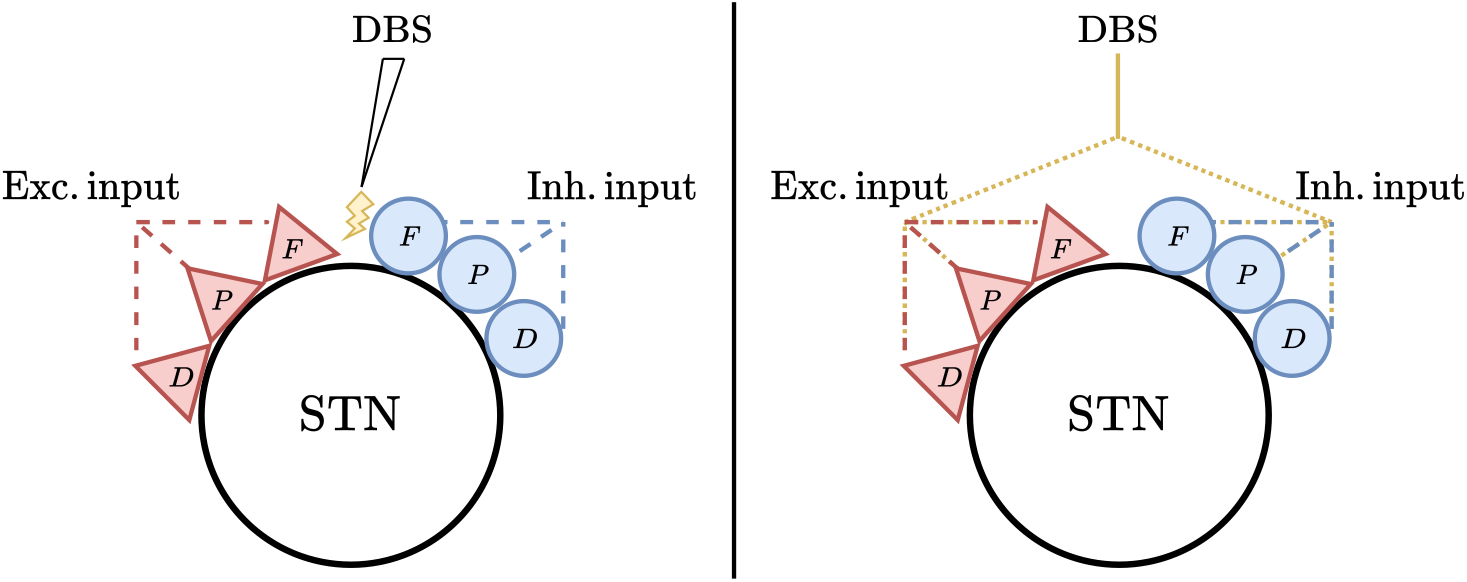
Conceptual framework to simulate the effect of DBS pulses on singular neuron responses: (left) schematic of DBS action on a single neuron target; (right) simulation of DBS action by simultaneous activation of all presynaptic inputs (F: facilitating; D: depressing; P: pseudo-linear)

In the right-hand side of Figure 2, DBS action is simulated by injecting artificial spikes into all presynaptic channels simultaneously. As a result, the overall STP effect, broken down into facilitating, depressing and pseudo-linear behaviours, combines the responses produced by presynaptic activity and the artificial DBS spikes, both excitatorily and inhibitorily.

#### Implementation

Spike input generation is integral to simulating neuronal dynamics on SpiNNaker. As such, the generation of artificial DBS spikes simply extends extends existing functionality known as a *spike source array* (S. Furber & Bogdan, 2020). The latter defines a population object with neuron-like entities that can emit spikes at predefined times during the simulation runtime. Similarly, we propose a *dbs spike soure array* which precomputes the spike times based on a frequency of stimulation (in Hz), a start time and an end time, all of which may be specified by the user.

In order to project the newly-created DBS-spike input population towards the target neuron along the different TM synapses, we reuse the previously defined TM synapse object (whether F, D, or P and whether excitatory or inhibitory) as well as the connector object (e.g., *one-to-one*). We note that this approach is equivalent to combining the DBS-spike input population with other spike input populations prior to connecting presynaptic and postsynaptic populations. This alternative formulation is used for the time being as a result of constraints imposed by the sPyNNaker environment (see Section 4 for additional details regarding its mitigation).

### 2.3 Leaky Integrate-and-fire Neuron

The target (STN) neuron is modelled as a leaky integrate-and-fire (LIF) with the following current-based dynamics (S. Furber & Bogdan, 2020):

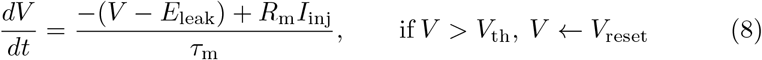

where *E*_leak_ = −60 mV is the membrane leak (resting) potential, *τ*_m_ = 10 ms is the membrane leak time constant and *R*_m_ = 1 MΩ is the membrane resistance. If the membrane potential exceeds the static threshold *V*_th_ = −40 mV, it is algorithmically reset to *V*_reset_ = −90 mV for the duration of a refractory period *τ*_r_ = 1 ms. The total current injection *I*_inj_ breaks down into

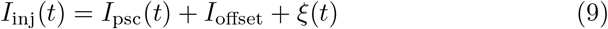

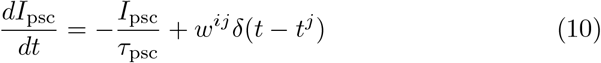

where *I*_offset_ = 0 nA denotes an intrinsic current (not considered in this case) and *ξ*(*t*) = 0 nA synaptic noise (while not implemented in this case, an Ornstein-Uhlenbeck process may be used). The dynamics of the postsynaptic current input *I*_syn_ are governed by the time constant *τ*_psc_ = 20 ms and the weighted contribution of incoming spikes (modelled as Dirac delta functions) from presynaptic neurons indexed by *j*. This is where neuron model and synapse model definitions overlap; given TM synapses between postsynaptic neuron *i* and presynaptic neuron *j, w*^*ij*^ is dynamic and evolves according to (7).

#### Implementation

In sPyNNaker, (8) corresponds to a *neuron model* component, whereas (10) corresponds to a *synapse type* component. On SpiNNaker, both the state of the neuron dynamics and the postsynaptic current are updated using exact integration (S. Furber & Bogdan, 2020). Given a simulation timestep *h* and the discrete-time index *k* = 0, …, *N*_sim_,

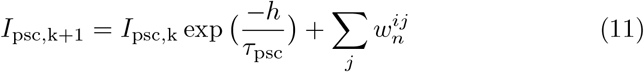

where the constant factor exp 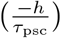 is precomputed in the corresponding Python interface and loaded onto the SpiNNaker hardware at the onset of simulation. Also, we recall the discrete TM weight update rule 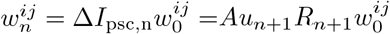 which is governed by the time interval between incoming spikes Δ*t*_*n*_ and ensures *I*_psc,k_ reflects changes in synaptic efficacy.

Similarly, the discrete membrane potential update is given by

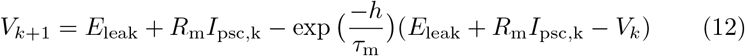

where the constant factor exp 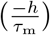 is once again precomputed.

To add neuron and synapse models to sPyNNaker, the desired components, existing or custom, need to be specified at compile time by referencing each component in the same Makefile. Given the close relationship between the synaptic dynamics and the neuron model, a new model is required to simulate STP with an LIF target neuron. Starting with the standard current-based LIF neuron model (8) (with exponential post-synaptic shaping and static threshold), we include the proposed TM timingand weightdependence rules as modified ST(D)P dynamics. Then, in order to support the facilitating, depressing and pseudo-linear TM types, both excitatorily and inhibitorily, we modify the exponential synapse type to support all six variations: EXC FAC, EXC DEP, EXC PSD, INH FAC, INH DEP, INH PSD (see `“NEURON”` section in Figure 1). The resulting Makefile is shown in Listing 2.3 to highlight the combination of existing functionality (directory references prefixed by `$(NEURON DIR)`) and the added functionality detailed in this work (prefixed by $(EXTRA SRC DIR)).

##### Listing 1

Makefile for the `IF_curr_exp_stdp_mad_tm_timing_tm_weight` model

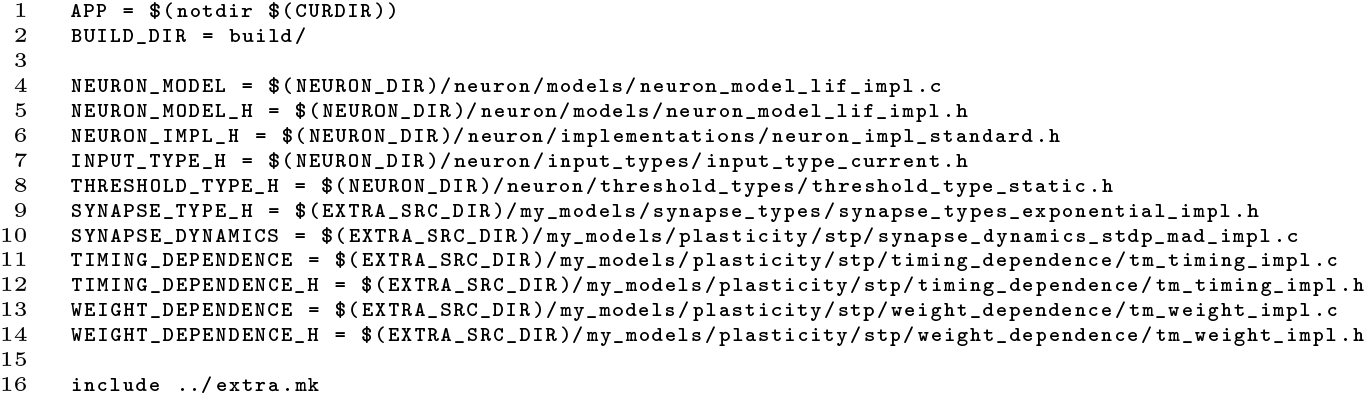

After writing the corresponding Python interface for the new exponential integrate-and-fire neuron model with support for TM synapses, STP dynamics may be simulated in PyNN using the structures previously associated with STDP dynamics as illustrated in the following section.

## 3 Results

The following results showcase various features of the proposed STP implementation based on the TM synapse model. The simulations consider the effect of different types of spiking input on the response of a single LIF target neuron.

### 3.1 Single Synapse

As mentioned previously, the TM implementations supports six synapse types; Figure 3 illustrates the membrane voltage and the postsynaptic current for three of these types, namely excitatory-facilitating (EXC FAC), excitatorydepressing (EXC DEP) and inhibitory-pseudo-linear synapses (INH DEP). The input consists of five spikes selected to mimic the STP examples in (M. Tsodyks & Wu, 2013).

**Figure 3.**
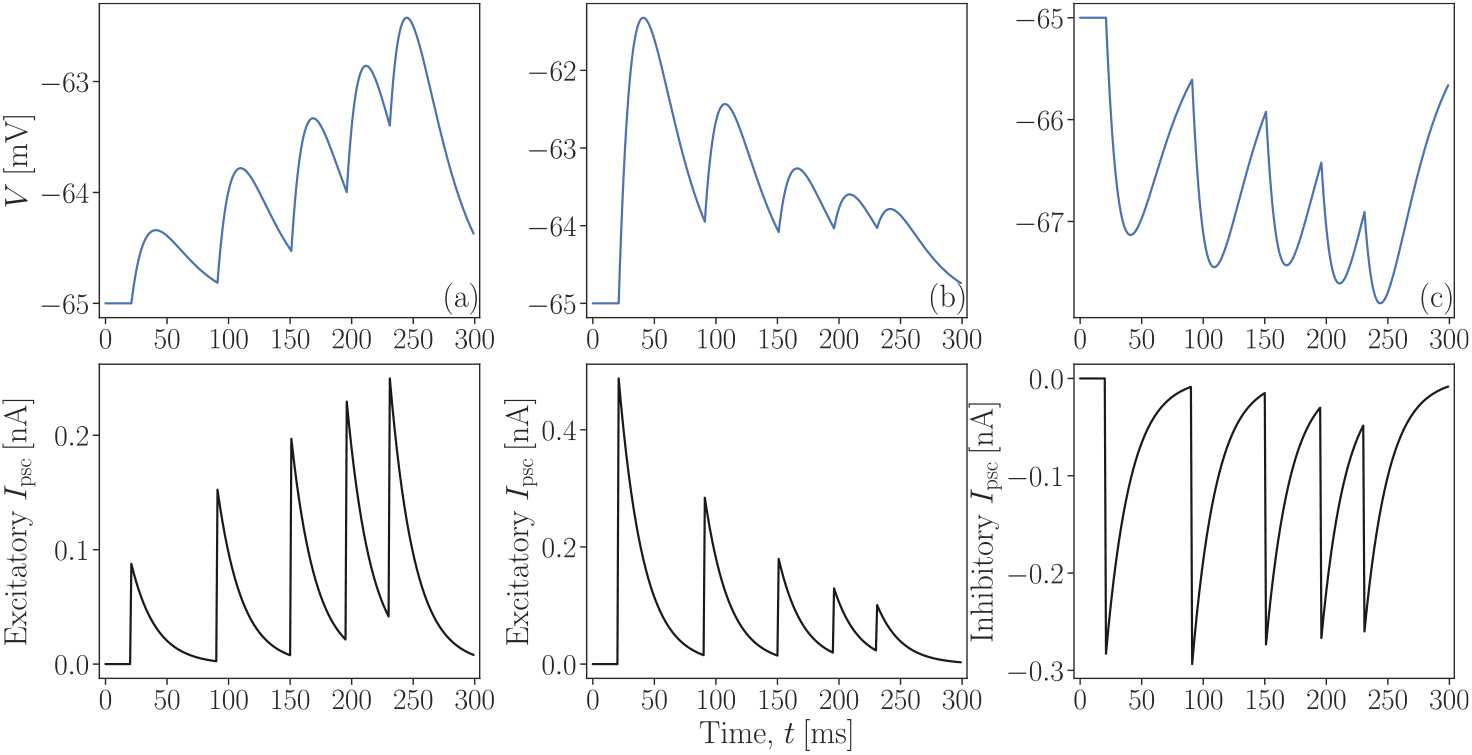
TM synapses connected to a single postsynaptic LIF neuron and subject to the input spike train *{*20, 90, 150, 195, 230*}* ms: (a) excitatory-facilitating synapse (*τ*_*f*_ = 670 ms, *τ*_d_ = 138 ms, U = 0.09); (b) excitatory-depressing synapse (*τ*_*f*_ = 17 ms, *τ*_d_ = 671 ms, U = 0.50); (c) inhibitory-pseudo-linear synapse (*τ*_*f*_ = 62 ms, *τ*_d_ = 144 ms, U = 0.29, *τ*_psc_ = 20 ms)

The facilitating and depressing effects are clearly represented in (a) and (b) respectively, while (c) depicts a combination of the two which is characteristic of pseudo-linear synapses. In the latter, the *I*_psc_ plot is negated to show that the input current inhibits the target neuron.

#### Hidden TM Dynamics

By default, only the membrane voltage and the excitatory and inhibitory postsynaptic currents (or the corresponding conductances in a conductance-based model) are returned by the SpiNNaker hardware at the end of the simulation. In studying STP with the TM model, it is beneficial to plot the evolution of the synaptic resources variables *u* and *R*. While steps can be taken in sPyNNaker to modify the number of recorded state variables, we have elected to retrieve their values from a log file for the time being. Figure 4 depicts the hidden dynamics of an excitatory-depressing synapse (middle plot) subject to noisy excitatory input spikes (top plot). The LIF membrane potential response is compared to the response obtained in NEST (via PyNN) under identical simulation settings (bottom plot).

**Figure 4.**
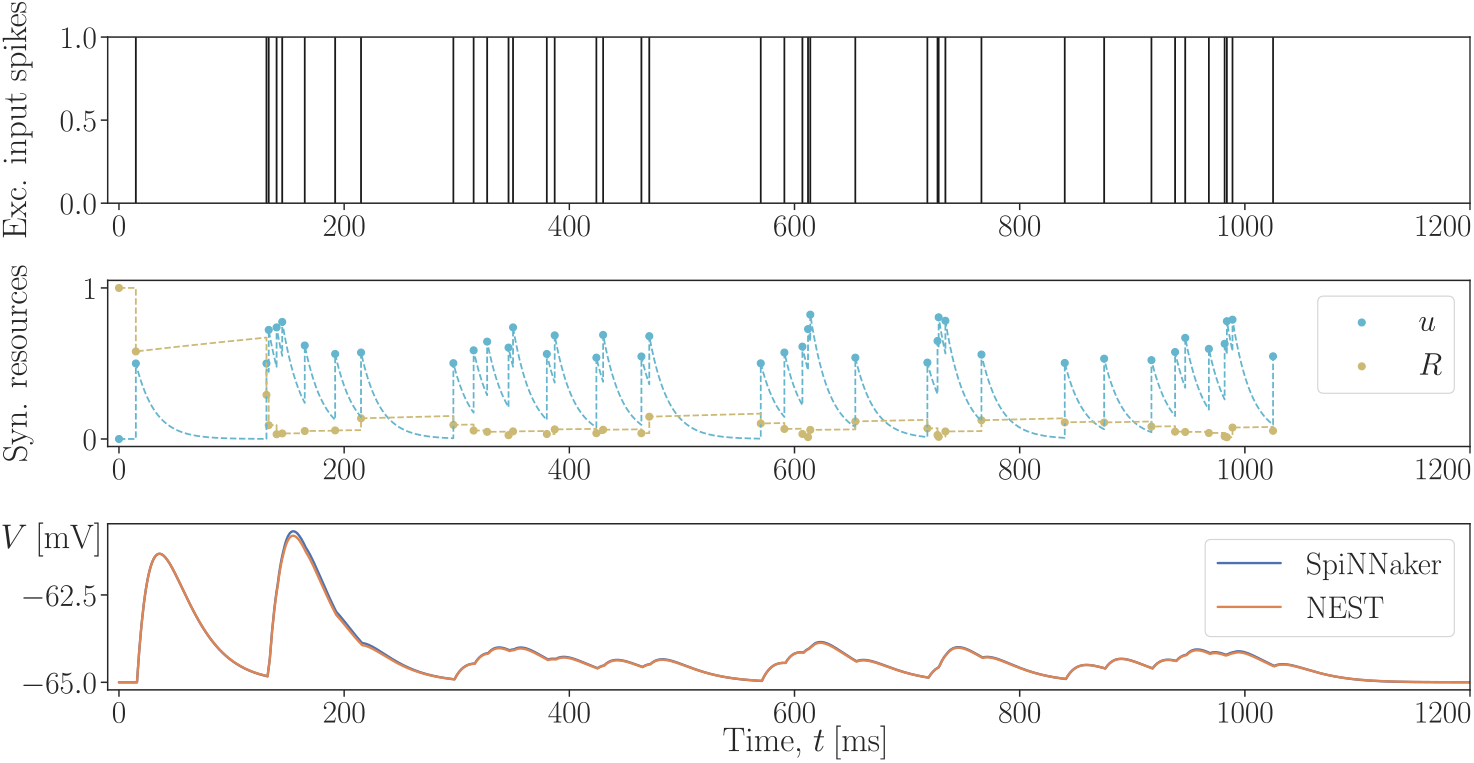
Excitatory-depressing TM synapse subject to noisy excitatory input spikes: the scatter points denoting the value of the synaptic resource variables *u* and *R* are recovered from the SpiNNaker output logs, while the exponential decay/growth interspike segments (dashed) are filled in during postprocessing (*τ*_*f*_ = 17 ms, *τ*_d_ = 671 ms, U = 0.50, *τ*_psc_ = 20 ms)

The slow recovery of the TM variable *R* in the middle plot above reflects the dominating depressive dynamics associated with the depletion of neurotransmitter resources. Despite an initial gain (caused by a rapid succession of incoming spikes right before the 200 ms mark), the postsynaptic neuron’s response is heavily suppressed as a result.

Slight inconsistencies in the membrane potential responses can be observed between the SpiNNaker and NEST backends, in particular when the input spike density is high. The source of the discrepancy will be discussed in Section 4. However, qualitatively, it appears the overall STP dynamics based on our proposed implementation for SpiNNaker are adequately captured.

#### Steady-State Behaviour

To quantitatively assess the extent to which our implementation agrees with the theoretical TM model, we evaluate the steady-state behaviour of the synapse as reviewed earlier (Markram et al., 1998). We make use of the proposed *dbs spike source array* to inject spikes into the target neuron at a desired frequency *f*_DBS_. Figure 5 compares the steady-state response of an excitatory-pseudo-linear (EXC PSD) synapse stimulated at a medium frequency of *f*_DBS_ = 50 Hz on the left to that of an inhibitory-depressing (INH DEP) synapse at a low frequency of *f*_DBS_ = 10 Hz on the right. On the right, the steady-state postsynaptic current *I*_psc,*∞*_ is negated to reflect an inhibiting input.

**Figure 5.**
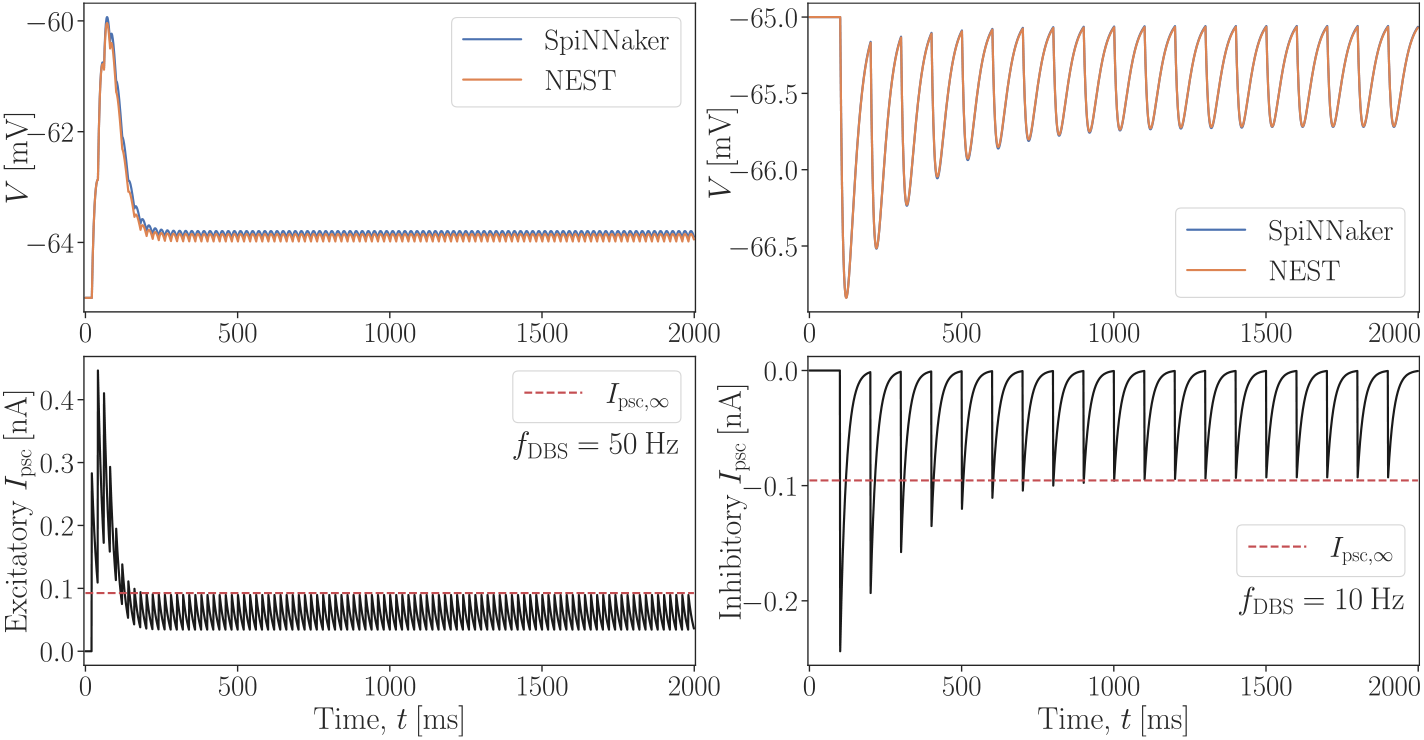
Steady-state behaviour of TM synapses subject to incoming spike trains of constant frequency *f*_DBS_: (left) excitatory-pseudo-linear synapse (*τ*_*f*_ = 326 ms, *τ*_d_ = 329 ms, U = 0.29); (right) inhibitory-depressing synapse (*τ*_*f*_ = 21 ms, *τ*_d_ = 706 ms, U = 0.25, *τ*_psc_ = 20 ms)

At first glance, the steady-state postsynaptic current *I*_psc,*∞*_ obtained from (6) coincides with the peak (negative peak) *I*_psc,peak_ of the excitatory (inhibitory) synaptic current simulated on SpiNNaker. Table 1 provides a quantitative assessment of the disparity.

**Table 1.**
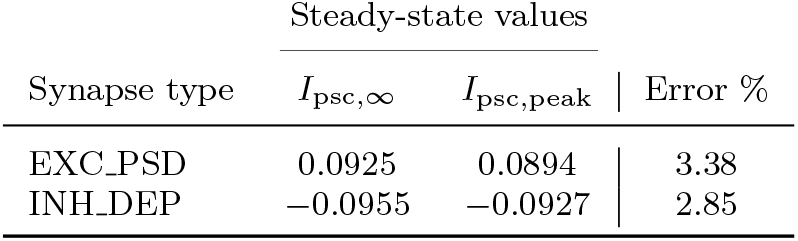
Steady-state comparison of the theoretical and simulated postsynaptic currents

| Synapse type | Steady-state values |  |  |
| --- | --- | --- | --- |
| | $I_{\text{psc},\infty}$ | $I_{\text{psc,peak}}$ | Error % |
| EXC_PSD | 0.0925 | 0.0894 | 3.38 |
| INH_DEP | -0.0955 | -0.0927 | 2.85 |

We note in passing that the discrepancy in membrane potential responses between NEST and SpiNNaker backends is once again appreciable at high frequency (Figure 5, top-left plot).

### 3.1.1 Multiple Synapses

Having demonstrated the dynamics of singular TM synapses, we now consider the simulation of combined STP effects as a result of multiple synapses converging to the same postsynaptic target. While not enabled by default on SpiNNaker, this functionality can be achieved by identifying the different TM synapse types in the Python simulation code and passing the information to the corresponding low-level TM implementation. For the time being, only one set of six TM synapse types is supported using this approach.

The following simulation considers three input populations, each with a different spike rate and each projecting to the same postsynaptic LIF neuron via TM synapses: the first synapse type is EXC FAC, the second EXC PSD and the third INH DEP. Figure 6 illustrates a superposition of STP effects as simultaneous incoming spikes converge to the target along different synapses.

**Figure 6.**
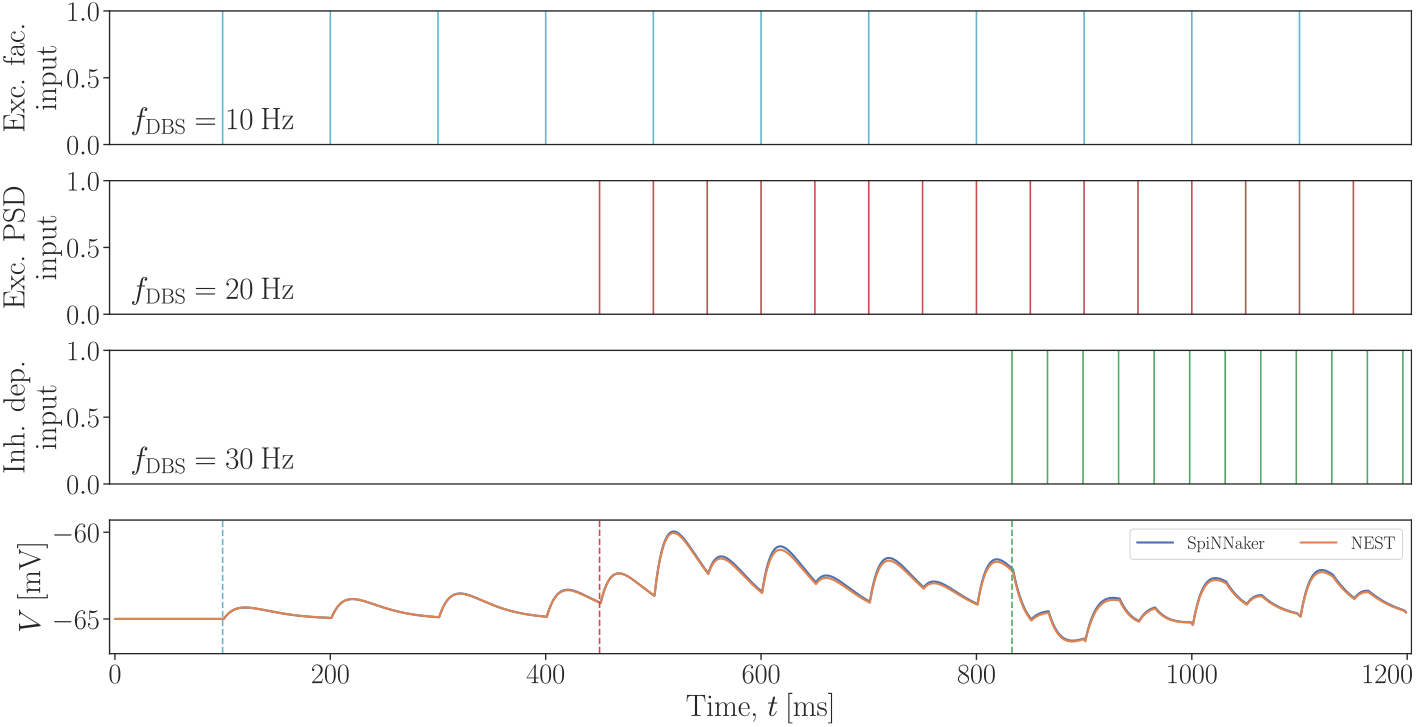
Superposition effect of multiple TM synapses connecting to the same postsynaptic neuron: (top) excitatory-facilitating synapse (*τ*_*f*_ = 670 ms, *τ*_d_ = 138 ms, U = 0.09); (middle) excitatory-pseudo-linear synapse (*τ*_*f*_ = 326ms, *τ*_d_ = 329ms, U = 0.29); (bottom) inhibitory-depressing synapse (*τ*_*f*_ = 21 ms, *τ*_d_ = 706 ms, U = 0.25, *τ*_psc_ = 20 ms)

The membrane potential response in the bottom plot is segmented to highlight the effects of respectively one, two and three dynamic synapses. Along the first two segments, the net effect is excitatory; however, the injection of inhibitory input at a higher frequency rapidly balances the overall postsynaptic response.

#### DBS Effect

The simulation in Figure 6 validates the superposition effect of multiple TM synapses in a simple scenario. We next test the functionality in a manner that is more relevant to the study of DBS effects. Specifically, we are interested in reproducing (and ultimately, extending) the results on DBS-induced STP advanced by Milosevic et al. (2021).

We recall the conceptual framework proposed in Figure 2, whereby DBS action is approximated by simultaneously activating all presynaptic inputs converging onto a target neuron. In order to simulate this framework on SpiNNaker, we consider two noisy spike sources, one excitatory (in red) and one inhibitory (in blue), which project to a LIF neuron via all six types of supported TM synapses. As per the framework, a third input population (in yellow) injects artificial DBS spikes into all six channels at a predefined frequency *f*_DBS_. Figure 7 illustrates the effect low-frequency stimulation has on the postsynaptic response, while Figure 8 illustrates the effect at high frequency (the parameters of the various synapses are referenced in the caption of previous simulation results). In each plot, the excitatory (respectively, inhibitory) postsynaptic current, combining the contributions of the F, D and P synapses, is plotted underneath the excitatory (respectively, inhibitory) spike inputs.

**Figure 7.**
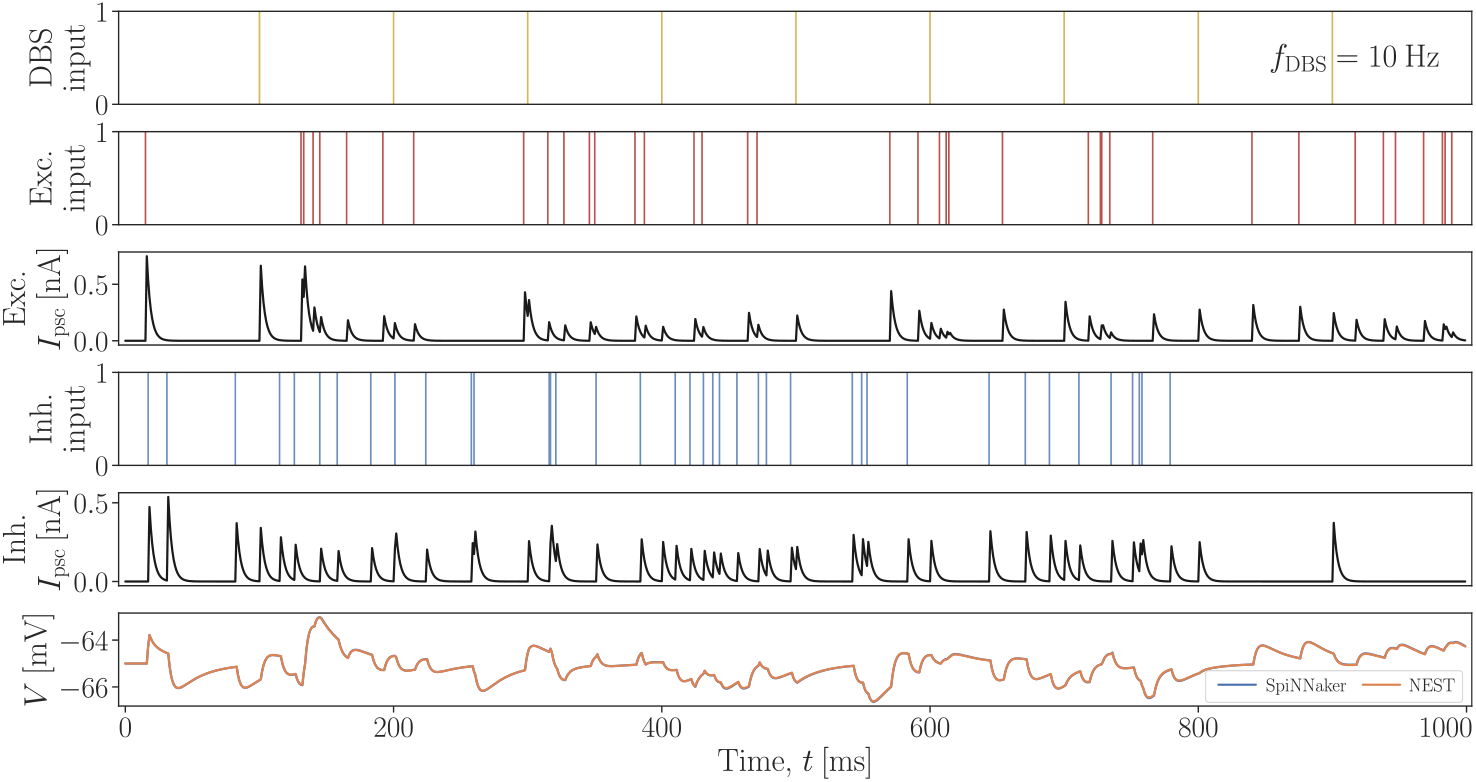
Simulation of low-frequency DBS impact on the response of a LIF neuron to which all six TM synapse types project (*f*_DBS_ = 10 Hz, *τ*_psc_ = 3 ms)

**Figure 8.**
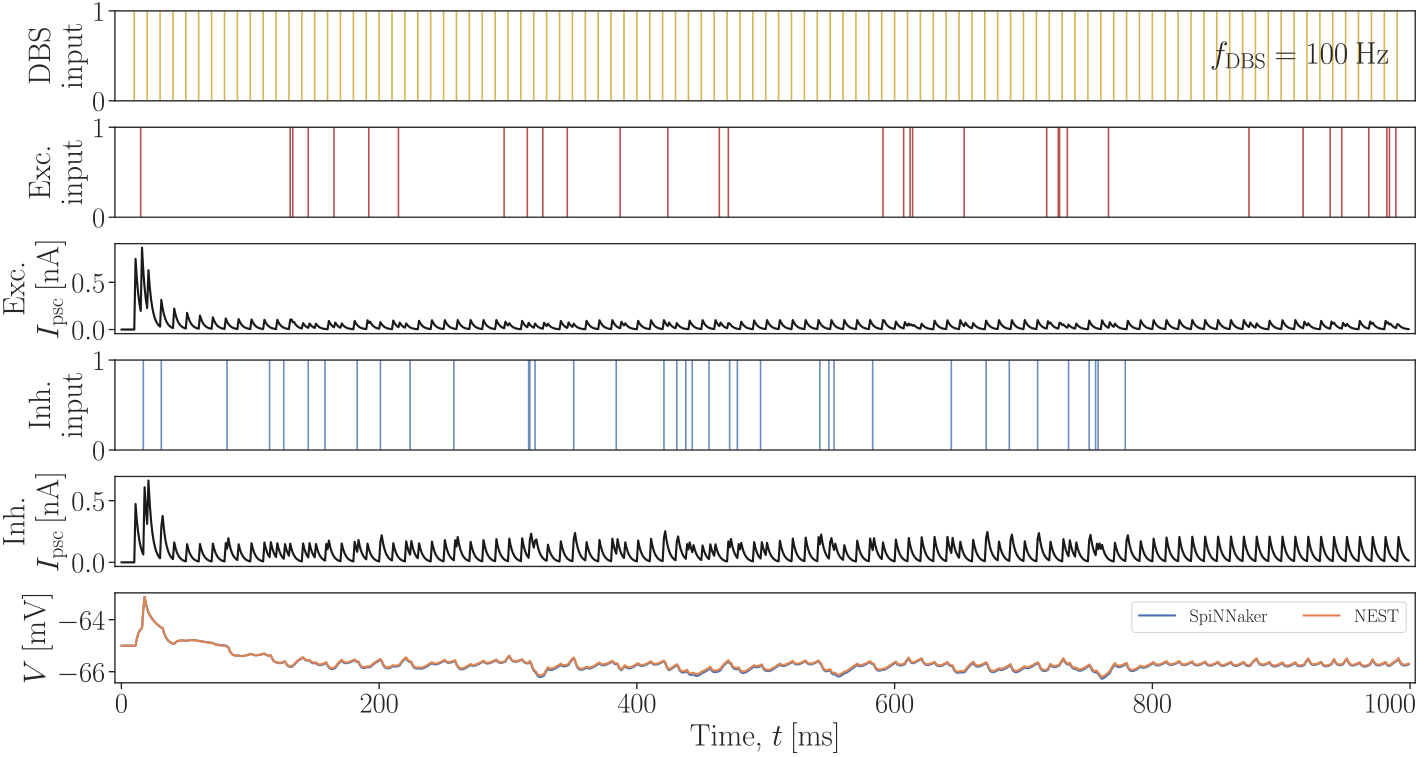
Simulation of high-frequency DBS impact on the response of a LIF neuron to which all six TM synapse types project (*f*_DBS_ = 100 Hz, *τ*_psc_ = 3 ms)

At low frequency, the excitatory and inhibitory postsynaptic currents, and the membrane potential response remain balanced and clear of any notable external influences. When combining the artificial spikes with the excitatory or inhibitory inputs, the resulting interspike intervals allow sufficient time for the various synapses to recover.

On the other hand, the high-frequency stimulation depicted in Figure 8 quickly saturates the synapses, both excitatorily and inhibitorily, as reflected by the combined postsynaptic currents (in black). The overall membrane potential response is heavily suppressed as a result. These simulations of the STP effect induced by DBS action agree with the findings advanced in (Milosevic et al., 2021).

## 4 Discussion

The proposed implementation of STP on SpiNNaker based on the TM model (Maass & Markram, 2002; Markram et al., 1998) produces encouraging result at the singular neuron level. Depending on parameterization, the effects of these new TM synapses may be studied individually or in combination with other synapse types, including other STP types. While early stage, the STP functionality already lends itself to studies of DBS impact on single neuron dynamics. The natural next step for this work consists in scaling up the simulations of STP from single neurons to populations, allowing studies of DBS effects to be extended to more realistic microcircuits.

Currently, there are several obstacles that limit such a change in scale. First, owing to the setup of our implementation, duplication of any one TM synapse type leads to incorrect behaviour. Consider a scenario similar to the one from Figure 8, where the DBS input source and the excitatory input population would project to the target using two different TM synapses of the same type (e.g., 2 EXC FAC synapses). While the time between presynaptic spikes Δ*t*_*n*_ would be respected independently for each synapse, the synaptic variables *u*_*n*_ and *R*_*n*_, and by extension the synaptic weight *w*^*ij*^, wound up being shared in the current implementation; as a result, the efficacy of each TM synapse does not reflect its presynaptic history. This issue significantly restricts the scope of STP simulations that can be considered with the current setup and it needs to be resolved to enable population-level scenarios.

Similarly, by choosing to extend the number of synapse types from the default two types (excitatory or inhibitory) to six types (excitatory or inhibitory, and, F, D or P), we offer a wider range of functionality for STP simulations. However, owing to the current setup of sPyNNaker, this solution effectively limits the total number of different TM synapses to six; that is, only one parameterization of each TM synapse type is supported at the time. While having one set of six TM synapse types may suit the simulation requirements of some, a less restrictive implementation is desired. We believe that, just like the six types are currently identified and passed to the low-level TM implementation, a more general solution would allow for the identification of unique TM synapses beyond those six. To do so, however, requires breaking from the excitatory or inhibitory, and, F, D or P structure currently offered by the *synapse type* components.

Both simulations of DBS-induced STP (Figures 7 and 8) include a comparison of membrane potential responses between SpiNNaker and NEST backends. Once again, slight discrepancies may be observed between the traces in instances of high stimulation. While it is likely that the fixed-point arithmetic on SpiNNaker may lead to some loss in precision, larger discrepancies could be observed as a result of changes in other simulation parameters (e.g., the synaptic time constant *τ*_psc_). This observation suggests that differences in implementation outside of the TM model might also be at play and in turn, these would be exacerbated when subject to frequent inputs. While qualitative correspondence between the two backends sufficed in this work, future work will look into the source of the disparity and the extent to which NEST can or should be used as a benchmark.

Thus far, we have limited our study of DBS and STP effects to subthreshold neuronal responses. Incorporating a bias current (or *offset* in sPyNNaker) in the target neuron effectively lifts the dynamics in a superthreshold regime. However, as a result of discrepancies in membrane potential responses between SpiNNaker and NEST backends, the corresponding postsynaptic spikes are emitted with small delays leading to increasing mismatch between the two traces. Again, matching postsynaptic spike timing between simulator backends may not be required. We expect the disparity at the singular neuron-level to have but a minor influence on the overall study of STP effects as larger simulations are considered.

Considering the variety of STDP rules currently supported by SpiNNaker, we envision additional implementations of STP in future extensions. While the phenomenological model proposed by (Markram et al., 1998) is commonly used to study STP dynamics, many extensions have been proposed since (BarrosoFlores et al., 2017; Hennig, 2013), some of which may prove more suitable for simulation on neuromorphic hardware. In particular, its nonlinear dependence on presynaptic spike history precludes any aggregation of spikes over multiple input sources, which, arguably, has the potential to simplify the implementation of STP on an embedded device. A recent study (Rossbroich, Trotter, Beninger, Tóth, & Naud, 2021) proposes a linear-nonlinear approach to STP where synaptic efficacy is modelled as a nonlinear readout of a convolution operation between the presynaptic spike train and an efficacy kernel. Given the ability to express the synaptic efficacy kernel in terms of linear basis functions, this approach shares similarities with the existing sPyNNaker implementation of postsynaptic current-shaping kernels. A future extension of this work will be dedicated to investigating the compatibility of this linear-nonlinear STP model with the SpiNNaker environment.

## Declarations

### Code availability

The Python and C code developed for this study follows the template repository provided by the sPyNNaker developers. It can be accessed publicly at https://github.com/nsbspl/sPyNNaker8NewModelTemplate.

## Acknowledgments

We thank Andrew Rowley from the University of Manchester SpiNNaker project for helpful assistance in developing this first TM model.

## Notes

### Competing Interest Statement

The authors have declared no competing interest.

## References

Barroso-Flores, J., Herrera-Valdez, M.A., Galarraga, E., Bargas, J. (2017). Models of Short-Term synaptic plasticity. Adv Exp Med Biol, 1015, 41–57.

Davies, M., Wild, A., Orchard, G., Sandamirskaya, Y., Guerra, G.A.F., Joshi, P., … Risbud, S.R. (2021). Advancing neuromorphic computing with loihi: A survey of results and outlook. Proceedings of the IEEE, 109 (5), 911–934.

Davison, A., Brüderle, D., Eppler, J., Kremkow, J., Muller, E., Pecevski, D., … Yger, P. (2009). Pynn: a common interface for neuronal network simulators. Frontiers in Neuroinformatics, 2.

Farokhniaee, A., & McIntyre, C.C. (2019). Theoretical principles of deep brain stimulation induced synaptic suppression. Brain Stimulation, 12 (6), 1402–1409.

Furber, S., & Bogdan, P. (2020). Spinnaker - a spiking neural network architecture. Boston-Delft: now publishers.

Furber, S.B., Galluppi, F., Temple, S., Plana, L.A. (2014). The spinnaker project. Proceedings of the IEEE, 102 (5), 652–665.

Gewaltig, M.-O., & Diesmann, M. (2007). Nest (neural simulation tool). Scholarpedia, 2 (4), 1430.

Ghadimi, A., Steiner, L.A., Popovic, M.R., Milosevic, L., Lankarany, M. (2021). Inferring deep brain stimulation induced short-term synaptic plasticity using novel dual optimization algorithm. bioRxiv.

Hennig, M.H. (2013, April). Theoretical models of synaptic short term plasticity. Front Comput Neurosci, 7, 45.

Lozano, A.M., & Lipsman, N. (2013). Probing and Regulating Dysfunctional Circuits Using Deep Brain Stimulation. Neuron, 77 (3), 406–424.

Lozano, A.M., Lipsman, N., Bergman, H., Brown, P., Chabardes, S., Chang, J.W., … Krauss, J.K. (2019, March). Deep brain stimulation: current challenges and future directions. Nature Reviews Neurology, 15 (3), 148– 160.

Maass, W., & Markram, H. (2002). Synapses as dynamic memory buffers. Neural Networks, 15 (2), 155–161.

Markram, H., Meier, K., Lippert, T., Grillner, S., Frackowiak, R., Dehaene, S., … Saria, A. (2011). Introducing the human brain project. Procedia Computer Science, 7, 39-42. (Proceedings of the 2nd European Future Technologies Conference and Exhibition 2011 (FET 11))

Markram, H., Wang, Y., Tsodyks, M. (1998). Differential signaling via the same axon of neocortical pyramidal neurons. Proceedings of the National Academy of Sciences, 95 (9), 5323–5328.

Meier, K. (2015). A mixed-signal universal neuromorphic computing system. 2015 ieee international electron devices meeting (iedm) (p. 4.6.1-4.6.4).

Milosevic, L., Kalia, S.K., Hodaie, M., Lozano, A.M., Popovic, M.R., Hutchison, W.D., Lankarany, M. (2021). A theoretical framework for the site-specific and frequency-dependent neuronal effects of deep brain stimulation. Brain Stimulation, 14 (4), 807–821.

Rhodes, O., Bogdan, P.A., Brenninkmeijer, C., Davidson, S., Fellows, D., Gait, A., … Furber, S.B. (2018). spynnaker: A software package for running pynn simulations on spinnaker. Frontiers in Neuroscience, 12.

Rossbroich, J., Trotter, D., Beninger, J., Tóth, K., Naud, R. (2021, 03). Linear-nonlinear cascades capture synaptic dynamics. PLOS Computational Biology, 17 (3), 1–27.

Sen-Bhattacharya, B., James, S., Rhodes, O., Sugiarto, I., Rowley, A., Stokes, A.B., … Furber, S.B. (2018). Building a spiking neural network model of the basal ganglia on spinnaker. IEEE Transactions on Cognitive and Developmental Systems, 10 (3), 823–836.

Tsodyks, M., & Wu, S. (2013). Short-term synaptic plasticity. Scholarpedia, 8 (10), 3153. (revision #182521)

Tsodyks, M.V., & Markram, H. (1997). The neural code between neocortical pyramidal neurons depends on neurotransmitter release probability. Proc Natl Acad Sci U S A, 94 (2), 719–723.

